# Decellularized biohybrid nerve promotes motor axon projections

**DOI:** 10.1101/2024.05.24.595727

**Authors:** Abijeet Singh Mehta, Sophia L. Zhang, Xinran Xie, Shreyaa Khanna, Joshua Tropp, Xudong Ji, Rachel Daso, Colin K. Franz, Sumanas W. Jordan, Jonathan Rivnay

## Abstract

Developing nerve grafts with intact mesostructures, superior conductivity, minimal immunogenicity, and improved tissue integration is essential for the treatment and restoration of neurological dysfunctions. A key factor is promoting directed axon growth into the grafts. To achieve this, we developed biohybrid nerves using decellularized rat sciatic nerve modified by in situ polymerization of poly(3,4-ethylenedioxythiophene) (PEDOT). We compared nine biohybrid nerves with varying polymerization conditions and cycles, selecting the best candidate through material characterization. Our results showed that a 1:1 ratio of FeCl3 oxidant to ethylenedioxythiophene (EDOT) monomer, cycled twice, provided superior conductivity (>0.2 mS/cm), mechanical alignment, intact mesostructures, and high compatibility with cells and blood. To test the biohybrid nerve’s effectiveness in promoting motor axon growth, we used human Spinal Cord Spheroids (hSCSs) from HUES 3 Hb9:GFP cells, with motor axons labeled with green fluorescent protein (GFP). Seeding hSCS onto one end of the conduit allowed motor axon outgrowth into the biohybrid nerve. Our construct effectively promoted directed motor axon growth, which improved significantly after seeding the grafts with Schwann cells. This study presents a promising approach for reconstructing axonal tracts in humans.

## 1. Introduction

Peripheral nerve injury (PNI) affects 200,000 Americans [1, 2].The main challenge in reversing the impact of this injury lies in the limited regenerative capacity of humans to rebuild and restore damaged neural circuits [3–5]. The damage to the nerve can be categorized into three distinct groups [6]. The most severe is third-degree nerve damage, which involves a complete transection or severing of the nerve causing a nerve gap [7]. The treatment method for peripheral nerve injuries (PNI) varies depending on the extent of damage [8]. The current standard practice for repairing nerve gap injuries involves using the patient’s own nerves through a process called autologous nerve autografting. However, this method has certain limitations, such as the need to harvest functional nerves from the patient, which can lead to complications at the donor site and limited availability of suitable nerves [9]. Nerve allografts, which are grafts from another individual, have also been extensively studied, but they require strong immunosuppressive treatment to prevent graft rejection and failure [10]. An alternative approach is the use of artificial nerve conduits, which must be biocompatible and capable of providing both physical and chemical signals for peripheral nerve repair. Ideally, these nerve conduits should have a biomimetic structure, possess sufficient mechanical properties to provide structural support and promote tissue integration, contain mesostructures to align the regenerating axons, allow for trophic support, conductivity, biodegradability, and flexibility [11]. However, the artificial nerve conduits reported in the literature do not meet all of these desired criteria nor accurately replicate natural conditions [8]. Natural decellularized nerve conduits are one promising solution [12].The striking advantage of decellularized nerve conduits is their potential to eliminate the need for autografts or allografts, thus overcoming the limitations associated with these conventional methods. The decellularized nerve, like autograft and unlike allograft/xenograft, induces minimal immunogenicity [13]. Additionally, decellularized nerve conduits offer similar topography and mechanical properties, provide guidance for axonal ingrowth, enhance tissue integration, can be customized for specific needs, allow for minimally invasive implantation, and are readily available [14].

The decellularization of the nerve can be achieved using various methods, such as chemical, physical, enzymatic, or a combination of these methods. The objective is to remove the cellular components, including nuclei, cytoplasm, and cellular debris, in order to eliminate potential immune-triggering elements while preserving the extracellular matrix (ECM). The ECM contains crucial structural proteins, growth factors, and signaling molecules necessary for tissue regeneration or other applications [15, 16]. However, it is important to note that the removal of primary cell types, such as neurons, glial cells (including astrocytes, oligodendrocytes, and Schwann cells), and endothelial cells, may also hinder the process of nerve regeneration. These cell types produce vital neurotrophic factors that can promote the growth of neurites from neurons [17]. Thus, there exists a tradeoff between the potential for inducing neurite outgrowth and the risk to mitigate immune reactions. Decellularization of the nerve can also reduce its conductivity, and it has been previously reported that conductive neural conduits promote outgrowth of neurites [18–21]. Therefore, in our study, we aimed to engineer a biohybrid nerve by combining a decellularized rat sciatic nerve with the conductive polymer poly(3,4- ethylenedioxythiophene) (PEDOT).

PEDOT is biocompatible [22, 23] and can facilitate cell migration and neurite outgrowth [24–27]. PEDOT and other conducting polymers have been extensively utilized in the field of tissue regeneration in recent years due to their unique combination of electrical conductivity and processability [28–31]. Previously, EDOT was polymerized *in-situ* within decellularized tissue utilizing ferric chloride (FeCl3) as an oxidant. The polymerization of PEDOT within decellularized tissue resulted in significantly higher electrical conductivities and it has been speculated that this biohybrid structure may facilitate peripheral nerve repair [32, 33]. However, there is a notable absence of a comprehensive study to validate the biophysical properties of such biohybrid nerves, optimized for deposition conditions, and to demonstrate their potential for tissue regeneration.

In this work, we investigated the biophysical properties and effectiveness of biohybrid nerve grafts, comprised of decellularized nerve and PEDOT. We tested the electrical, mechanical, and biocompatibility properties of constructs with various PEDOT polymerization conditions, changing oxidant to monomer ratio and polymerization cycle (number of rounds of polymerization). To compensate for the loss of neurotrophic factors due to complete decellularization of nerve graft we also seeded biohybrid nerve with human Schwann cells (SCs). We focused on investigating targeted growth of spinal motor axons into biohybrid nerves using hSCS, which are three-dimensional cell culture models derived from human pluripotent stem cells (hPSCs). These spheroids mimic the organization and functionality of the human spinal cord [34]. We derived hSCS from HUES 3 Hb9:GFP cells to utilize green fluorescent protein (GFP) to label the motor axons. By using GFP labeling, we ensured reliable and clear observations, which assisted in drawing comparison between motor axons projections into the biohybrid nerves and the decellularized peripheral nerve donor. Our findings demonstrated that our optimized biohybrid constructs successfully promoted the ingrowth of spinal motor axons when compared to pristine decellularized nerve (PDN).

## 2. Results

### 2.1. Engineering conductive biohybrid nerve conduits

To harness the advantages of allografts, such as similar structure, better support for axons, convenience of immediate use, avoidance of donor defects, and a seemingly endless supply, we harvested rat sciatic nerves from the hind limb in a sterile environment **(Figure 1A,B)**. In order to minimize the risk of immune responses, we decellularized the nerves by removing neurons, glial cells, and endothelial cells, as explained in the materials and methods section **(Figure 1C)**. We preserved the nerve architecture and guidance cues provided by extracellular matrix (ECM) components [35, 36] and to compensate for the loss of glial cells, which affects the physical and chemical cues as well as the conduction properties, we introduced the conductive polymer PEDOT to the nerve **(Figure 1D).** The decellularized nerves were cut into 5mm pieces and then treated with PEDOT through in situ polymerization. PEDOT is a biocompatible conductive polymer, it also offers sufficient cues to facilitate cell migration and the growth of neurites [24–27, 37].To examine the influence of polymerization conditions on the conductivity of the biohybrid nerves, we engineered four composites (DNP1:0.5C1, DNP1:1C1, DNP1:2C1, and DNP1:4C1) by varying the concentration of ferric chloride (FeCl3) in relation to a fixed concentration of 1M monomer EDOT **(Figure 1E)**. The FeCl3 concentration ranged from 0.5M to 4M. Subsequently, to enhance the conductive properties of the biohybrid nerves, we subjected one of the aforementioned composites (DNP1:1C1) to additional cycles of *in-situ* polymerization, ranging from 2 to 6 **(Figure 1F)**. This process resulted in the creation of five additional biohybrid nerves: DNP1:1C2, DNP1:1C3, DNP1:1C4, DNP1:1C5, and DNP1:1C6. All nine composites were fabricated using the same procedure described in the materials and methods section.

**Figure 1.**
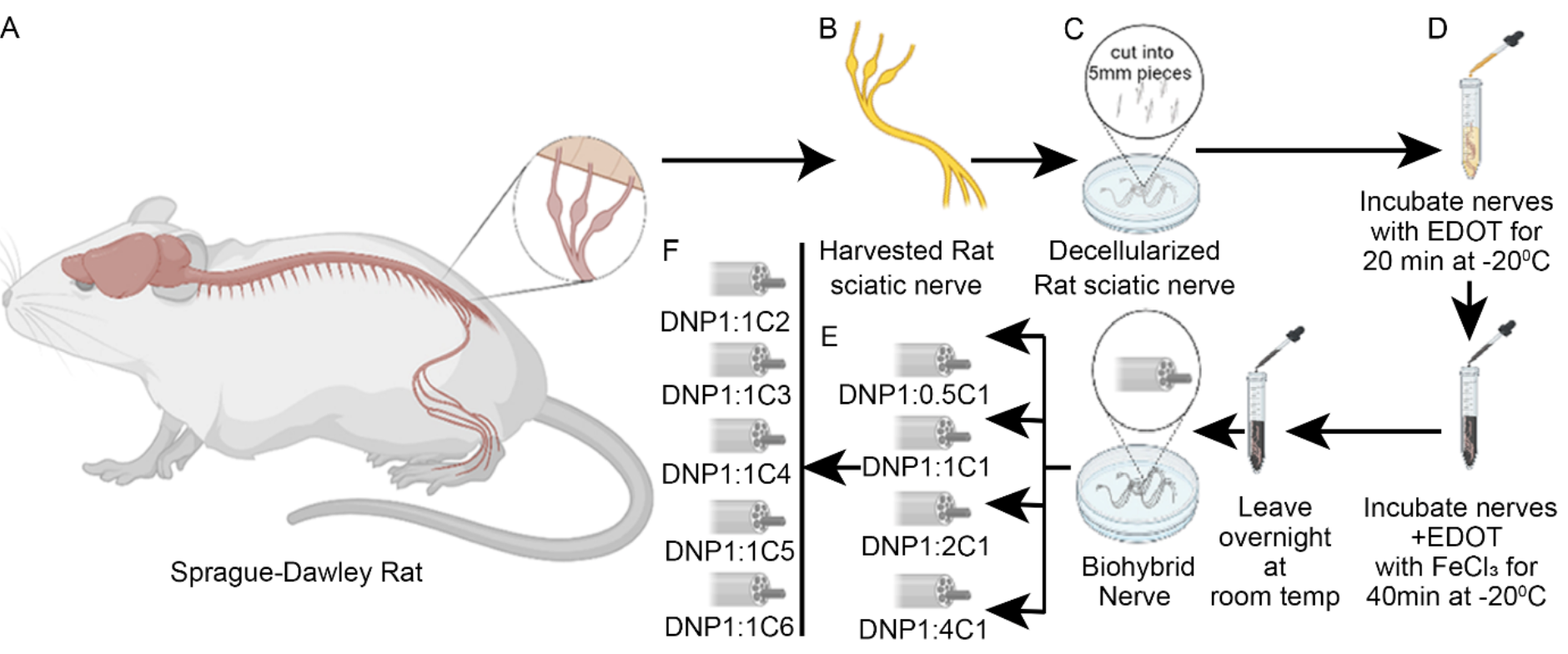
Engineering conductive biohybrid nerves. (A) Sciatic nerves were harvested from euthanized female Sprague Dawley rats under aseptic conditions. (B) The isolated sciatic nerves were then cleared of peripheral fat and connective tissue. (C) This was followed by a decellularization process, involving a freeze and thaw cycle in PBS, treatment with detergents, and sonication. **T**he nerves were then cut into 5mm pieces and stored at room temperature. (D) The conductive polymer PEDOT was integrated into the decellularized nerves through in-situ polymerization. (E) By varying the molar concentration of FeCl3 respective to the molar concentration of EDOT, four different biohybrid nerves were generated: DNP1:0.5C1, DNP1:1C1, DNP1:2C1, and DNP1:4C1. The biohybrid nerve DNP1:1C1 underwent subsequent cycles of polymerization, resulting in the fabrication of (F) five more biohybrid nerves: DNP1:1C2, DNP1:1C3, DNP1:1C4, DNP1:1C5, DNP1:1C6. This image was created with BioRender.com.

Next, to verify the successful polymerization of EDOT within the nerve composites we conducted FTIR analysis **(Figure S1A).** The FTIR spectra exhibit distinct bands corresponding to collagen. These bands appear at approximately 3400 cm^-1^ and 1625 cm^-1^, respectively (referred to as amide A and amide I). Such bands are characteristic of collagen-based materials. The amide I band represents the stretching of C=O bonds, which is influenced by the hydrogen bonding within the secondary structure. Therefore, the absence of any noticeable shift in the amide I band suggests that the secondary structure remained largely unaffected after multiple cycles of polymerization [38–40]. Following the polymerization of EDOT, new bands emerge at around 1520 cm^-1^ and 1320 cm^-1^. These bands correspond to the stretching vibrations of C=C and C-C bonds in the thiophene ring, respectively [41–44]. Additionally, characteristic peaks at approximately 980 cm^-1^, 830 cm^-1^, and 680 cm^-1^ appear, which correspond to the stretching vibrations of C-S-C bonds. Furthermore, the absence of C-H stretching bands in the range of 3100 cm-^1^ to 3050 cm^-1^ suggests that all thiophene rings within the composite are in the polymeric state rather than the monomeric state. These results suggest the successful polymerization of EDOT within the composite, an increased incorporation of PEDOT with each successive cycle, and an unaffected secondary structure of the collagen within the nerve composite.

Additionally, we employed X-ray Fluorescence (XRF) to quantify the relative sulfur concentrations in the biohybrid nerve samples relative to PDN. The data clearly demonstrates a gradual increase in relative sulfur with each cycle of PEDOT polymerization **(Figure S1B)**. Since each incorporated EDOT structural unit within the PEDOT polymer contains a sulfur atom, the relative sulfur content can serve as an indicator of the amount of PEDOT in the nerve and therefore data suggests that increase in the polymerization cycle as expected lead to an increase in the amount of PEDOT within the sample.

### 2.2. Electrical characterization of biohybrid nerves

The objective of engineering a biohybrid nerve is to improve its conductivity. To investigate its electrical properties, we conducted Electrochemical Impedance Spectroscopy (EIS). The results are presented through a Bode plot, which displays the magnitude of impedance (logarithmic scale) and phase angle in relation to frequency (logarithmic scale) **(Figure 2)** and Nyquist plot (Zimag versus ZReal) **(Figure S2)**. Both plots show that the electrode with nerve samples exhibit both resistive and capacitive behavior, including resistive behavior at higher frequencies, and a more capacitive behavior at lower frequencies for non-biohybrid nerves (PDN and PnDN) whereas, biohybrid nerves show resistive behavior at both higher and lower frequencies. Consequently, we constructed an equivalent circuit model to characterize the electric properties of nerve samples, which comprised the following components: R1(Q1||(R2Q2)) for non-biohybrid nerve and R1(Q1||R2)(Q2||R3) for biohybrid nerves **(Figure 2B)**. The first component, R1 represents un-compensatory resistance, R2Q1 represent bulk nerve-liquid interface and R3Q2 represent PEDOT coating-liquid interface. The equivalent circuit closely fitted the acquired data with chi^2^ value less than 0.05.

**Figure 2.**
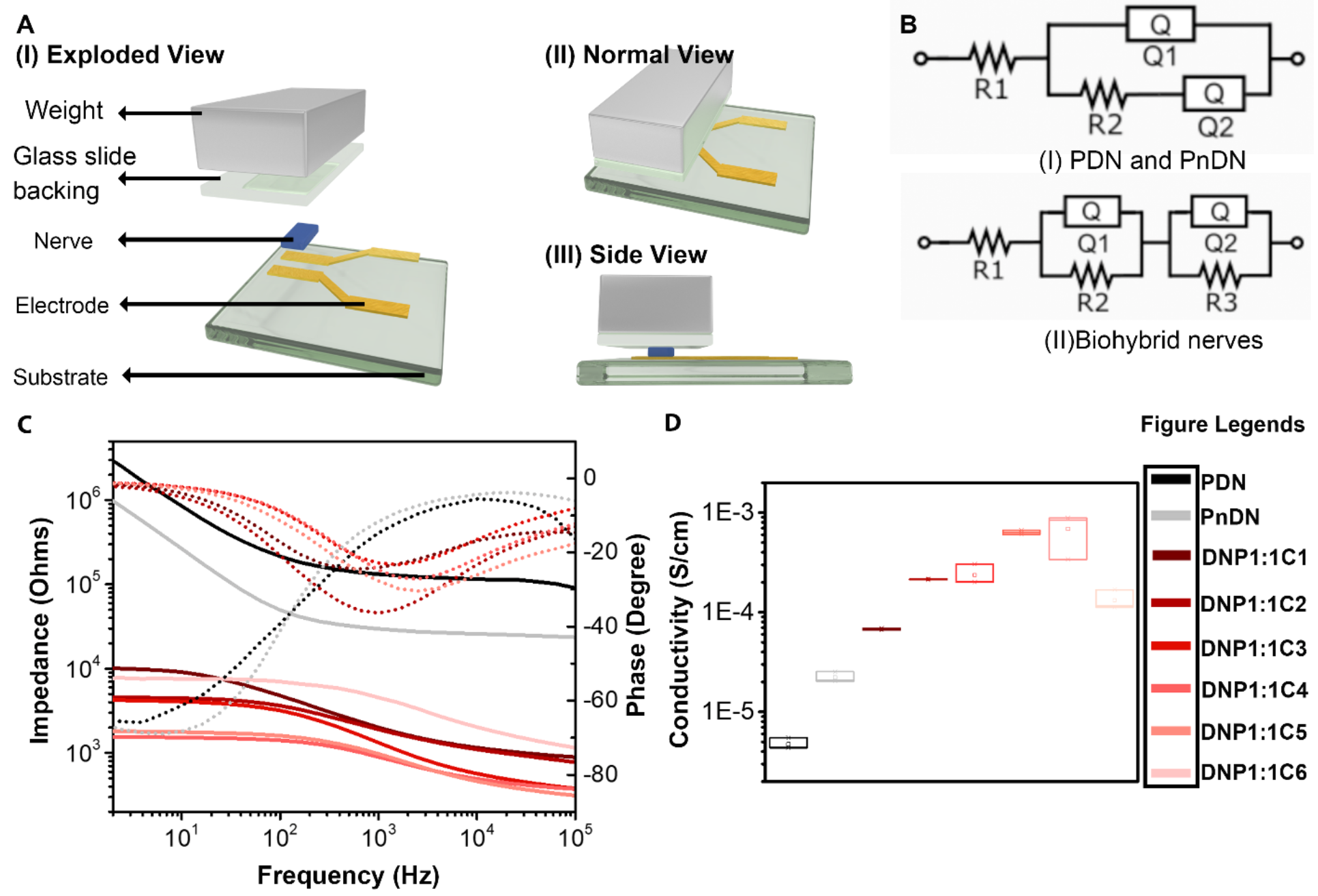
Evaluating electrical properties of biohybrid nerves. (A) The setup used to measure the conductivity of the biohybrid nerve consists of a glass substrate coated with gold, creating a 1mm gap. This gap is bridged by the nerve sample to complete the circuit. One more glass slide attached to the fixed weight is placed on top of the nerve to improve contact between nerve and gold electrode (B) The EIS technique is employed to measure the electrical properties, and the acquired data is fitted to R1(Q1**||**(R2Q2)) and R1(Q1||R2)(Q2||R3) circuit model. (C) The bode plot displays the impedance magnitude (solid lines) and phase angle (dotted lines) of pristine decellularized nerve (PDN), pristine non-decellularized nerve (PnDN), and biohybrid nerves with different cycles of polymerization (DNP1:1C1; DNP1:1C2; DNP1:1C3; DNP1:1C4; DNP1:1C5; DNP1:1C6). (D) The conductivity value for the biohybrid nerves with varying cycles of polymerization (DNP1:1C1; DNP1:1C2; DNP1:1C3; DNP1:1C4; DNP1:1C5; DNP1:1C6) is calculated and compared to PDN. Quantified as mean ± SD, n = 3 nerve samples in each group, *P < .05, **P < .01, ***P < .001, ns is non-significant.

The R2 and R3 value obtained from the circuit fit is used to calculate the conductivity of the nerve samples **(Figure 2D and Table S2)**. This is done using the formula: Conductivity (S) = Thickness (T) / (Charge Transfer (RCT) × Area (A)). Here RCT= R2+R3 [18]. The conductivity of the non-decellularized autograft (PnDN), in its original state, is measured to be 0.022 mS cm^-1^. However, after undergoing decellularization, the conductivity of the autograft (PDN) significantly decreases to 0.004 mS cm^-1^. This decrease could be attributed to the loss of conductive cellular components during the decellularization process. We then polymerized EDOT within the decellularized nerve. Initially, we investigated the effect of different polymerization conditions on the conductivity of the resulting biohybrids. Specifically, we varied the ratio of the catalyst ferric chloride (FeCl3) to the monomer EDOT. Among the investigated composites, we found that the biohybrid (DNP1:1C1) formed under a 1:1 molar ratio condition exhibited the highest conductivity of 0.067 mS cm^-1^.

It is important to note that the biohybrid nerves discussed above exhibited higher conductivity than both the autografts, PnDN, and PDN. However, we aimed to further enhance the conductivity. Therefore, we subjected the biohybrid nerve DNP1:1 to additional cycles of in situ polymerization ranging from 2 to 6 (DNP1:1C2, DNP1:1C3, DNP1:1C4, DNP1:1C5, DNP1:1C6). At cycle 2, DNP1:1C2 showed a dramatic increase in conductivity compared to DNP1:1 (DNP1:1C2 = 0.214 mS cm^-1^). Increasing the number of cycles to 3 and 4 resulted in similar trends in conductivity. The conductivity values for biohybrid nerve DNP1:1C3 is 0.236 mS cm^-1^ and for DNP1:1C4 is 0.636 mS cm^-1^. After adding one more cycle, the conductivity reached its peak value with DNP1:1C5 at 0.690 mS cm^-1^, and it started decreasing again for the subsequent cycle, DNP1:1C6, with a conductivity of 0.132 mS cm^-1^ **(Figure 2D)**.

The overall electrical/electrochemical characterization of biohybrid nerves demonstrated that polymerizing nerves with PEDOT enhances their conductivity. Additionally, subjecting the best polymerization condition (DNP1:1C1) to more polymerization cycles further enhances the conductivity, with the highest conductivity achieved by the biohybrid nerve DNP1:1C5= 0.690 mS cm^-1^.

### 2.3. Biophysical and biochemical characterization of biohybrid nerves

After evaluating the electrical properties of the biohybrid nerves, our goal was to examine their biophysical characteristics. This involved studying their mechanical properties, assessing the mesostructures of biohybrid nerves in comparison to pristine conditions, and quantifying the total amount of residual iron (Fe) in each biohybrid nerve.

We investigated the mechanical properties, specifically the stiffness, of the biohybrid nerves by calculating their compressive Young’s modulus **(Figure 3A,B)**. First, we assessed the effect of polymerization conditions on the Young’s modulus of the biohybrid nerves. The mean Young’s modulus values obtained for DNP1:0.5C1, DNP1:1C1, DNP1:2C1, and DNP1:4C1 are 43.01 kPa, 46.85 kPa, 56.44 kPa, and 67.78 kPa, respectively. The data clearly indicates that the stiffness of biohybrid nerves increases with an increase in the concentration of FeCl3 relative to the monomer EDOT. However, the change in Young’s modulus for all the aforementioned biohybrid nerves is not significantly different from that of the pristine nerve (PDN), which has a Young’s modulus value of 32.94 kPa **(Figure 3A).** This suggests that polymerizing the nerves intimately with PEDOT does not substantially affect their stiffness. Next, we quantified the change in Young’s modulus resulting from more than one layer of PEDOT polymerization of the biohybrid nerve DNP1:1C1. The mean Young’s modulus values obtained for DNP1:1C2, DNP1:1C3, DNP1:1C4, DNP1:1C5, and DNP1:1C6 are 88.39 kPa, 96.29 kPa, 219.24 kPa, 239.34 kPa, and 286.53 kPa, respectively. The stiffness of the biohybrid nerves significantly increases with each additional cycle from 2 to 6. However, two layers of PEDOT (DNP1:1C2) does not significantly change the stiffness compared to PDN (88.39 kPa vs. 32.94 kPa) **(Figure 3B)**.

**Figure 3:**
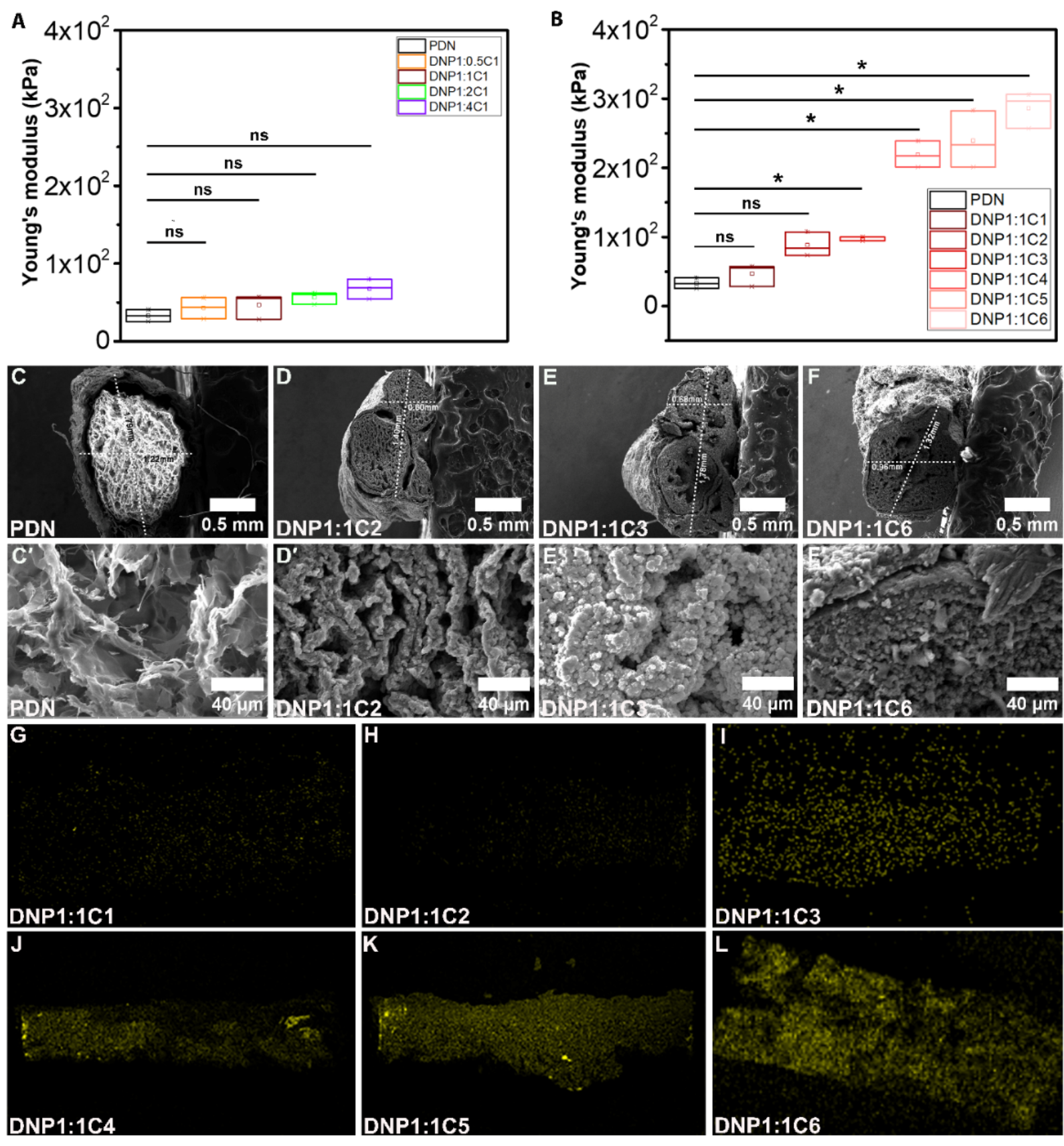
Evaluation of Mechanical properties, Nerve Topology and residual Fe content after in-situ polymerization with PEDOT. The Young’s modulus value for the biohybrid nerves with (A) changing concentrations of FeCl3 (DNP1:0.5C1; DNP1:1C1; DNP1:2C1; DNP1:4C1) and (B) varying cycles of polymerization (DNP1:1C1; DNP1:1C2; DNP1:1C3; DNP1:1C4; DNP1:1C5; DNP1:1C6) is calculated and compared to PDN. (C- F) Scanning electron microscopy images of biohybrid nerves (C,C′) PDN (D,D′) DNP1:1C2 (E,E′) DNP1:1C3 (F,F′) DNP1:1C6. (G-L) Energy-dispersive X-ray Spectroscopy (EDS) mapping of Fe within (G) DNP1:1C1 (H) DNP1:1C2 (I) DNP1:1C3 (J) DNP1:1C4 (K) DNP1:1C5 (L) DNP1:1C6. Scale bar = 0.5 mm(A-D) and 40 μm(A′-D′). Quantification is done as mean ± SD, n = 3 nerve samples in each group, *P < .05, **P < .001, ns is non-significant.

We used scanning electron microscopy (SEM) to examine the changes in the structure of biohybrid nerves before and after polymerization with PEDOT **(Figure 3C,C′-F,F′ and Figure S3A,A′, A ′′-J,J′, J ′′)**. The top view micrograph of PDN clearly reveals fibrillar structures on the nerve membrane **(Figure S3A,A′, A ′′)**, which are likely due to collagen fibers [45, 46]. There are no apparent changes in the fibrillar structures of biohybrid nerves DNP1:0.5 C1, DNP1:1C1, DNP1:2C1, and DNP1:4C1, despite variations in polymerization conditions and increasing concentrations of FeCl3 in relation to the monomer EDOT **(Figure S3B,B′,B′′-E,E′,E′′)**. As subsequent PEDOT are added, the fibrous structures gradually become thicker and eventually become concealed within the PEDOT layers. In the cases of DNP1:1C5 and DNP1:1C6, the fibrous structures are no longer discernible **(Figure S3F,F′,F′′-J,J′,J′′)**. However, up until the third round of PEDOT polymerization (DNP1:1C2, DNP1:1C3), the biohybrid nerves still clearly exhibit visible fibrous structures **(Figure S3F,F′,F′′,G,G′,G′′)**, which appear to be coated by globular particles which are likely PEDOT [47].To compare the changes in fascicular structures resulting from the application of multiple layers of PEDOT, we took side view images of the pristine nerve PDN and biohybrid nerves DNP1:1C2, DNP1:1C3, and DNP1:1C6 **(Figure 3C,C′-F,F′)**. The micrographs clearly demonstrate that the fascicular openings become densely packed with PEDOT in the case of the biohybrid nerve DNP1:1C6, which has undergone the maximum cycles of polymerization with PEDOT **(Figure 3F,F′)**. However, the nerve with two coatings of PEDOT (DNP1:1C2) retains visible openings and intact fascicular structures when compared with PDN **(Figure 3D,D′)**. Similarly, in DNP1:1C3, micropores are still visible but the mesostructures start to deteriorate **(Figure 3E,E′).**

Next, it was crucial to measure the remaining iron content in the biohybrid nerves. To achieve this, we used energy-dispersive X-ray spectroscopy (EDS) mapping and X-ray Fluorescence (XRF). The EDS micrographs provided clear evidence of residual iron on the nerve increase as the polymerization cycle with PEDOT progressed **(Figure 3G-L)**. To thoroughly investigate the iron content in the biohybrid nerves, we crushed nerve samples into powder and performed XRF analysis. The results indicate that both the pristine nerve PDN and the biohybrid nerve with a single polymerization cycle (DNP1:1C1) contained very little iron, with values below the limit of detection. After the first cycle, each subsequent cycle showed a slight increase in residual iron. The samples DNP1:1C2, DNP1:1C3, DNP1:1C4, DNP1:1C5, and DNP1:1C6 had relatively similar iron concentrations **(Figure S3K)**. This suggests that after the first cycle, iron becomes trapped within the sample during subsequent cycles, but the amount ranges from 0.5ppm to 29ppm, which is minimal and unlikely to have any harmful effects on animal health [48–51].

The overall analysis of the materials showed that the biohybrid nerve DNP1:1C2 exhibits minimal changes in stiffness. It maintains its mesostructures intact, and the concentration of residual Fe is low, posing no significant concerns for animal health.

### 2.4. Cytocompatibility and Hemocompatibility of biohybrid nerves

It is crucial to assess the compatibility of biohybrid nerves with cells (cytocompatibility) and blood (hemocompatibility) before determining their potential for reconstructing axonal tracts. The ISO 10993-5 standards recommend using mouse fibroblast cells (L929) to evaluate the cytocompatibility of biomaterials after their application and extraction [52]. In this study, the cytotoxicity of biohybrid nerves is evaluated using the MTT assay after culturing them with L929 cells for 48 hours and 72 hours **(Figure 4A,B)**. The results show that the cell viability percentage of the pristine nerve PDN is the highest, with 91.89% cell viability at 48 hours and 90.65% at 72 hours. However, the cell viability gradually decreased for the biohybrid nerves as the cycles of polymerization increased. Among the biohybrid nerves, the lowest cell viability was observed for biohybrid nerve DNP1:1 C6, with 78.37% at 48 hours and 76.29% at 72 hours. The biohybrid nerve of interest (DNP1:1C2), which exhibited excellent electrical, mechanical, and morphological characteristics, had a cell viability of 86.93% and 84.03% after 48 hours and 72 hours of cell cultivation, respectively. According to the ISO 10993-5 standards, any biomaterial with a cell viability percentage higher than 70% is considered cytocompatible [53–55]. Our data clearly demonstrate that both the pristine nerve (PDN) and all the biohybrid nerves are highly compatible with mammalian cells, as their cell viability percentages comfortably exceed 70% after 48 hours and 72 hours of cell cultivation **(Figure 4A,B)**. To further validate these findings and evaluate cell cytotoxicity in direct contact with the biohybrid nerves, we conducted a Live/Dead assay on the pristine nerve (PDN) and selected biohybrid nerves (DNP1:1C1, DNP1:1C2, DNP1:1C3) for testing. The images clearly demonstrate that all the tested nerve samples exhibited minimal cytotoxicity **(Figure 4C,C′,C′′,C′′′-F,F′,F′′,F′′′)**. The cell viability percentages for the pristine nerve PDN and biohybrid nerves DNP1:1C1, DNP1:1C2, DNP1:1C3 are 91.51%, 91.16%, 89.70%, and 87.30%, respectively **(Figure 4G)**. Additionally, the cells demonstrated intimate adherence to the material, as evident from their flattened morphology **(Figure 4C,C′,C′′,C′′′-F,F′,F′′,F′′′)**. However, there were clear differences in the overall number of L929 cells spreading and adhering to the biohybrid nerves, with the least attachment observed in DNP1:1C3 **(Figure 4F,F′,F′′,F′′′)**. This difference may be attributed to changes in film roughness due to an increased number of polymerization cycles.

**Figure 4.**
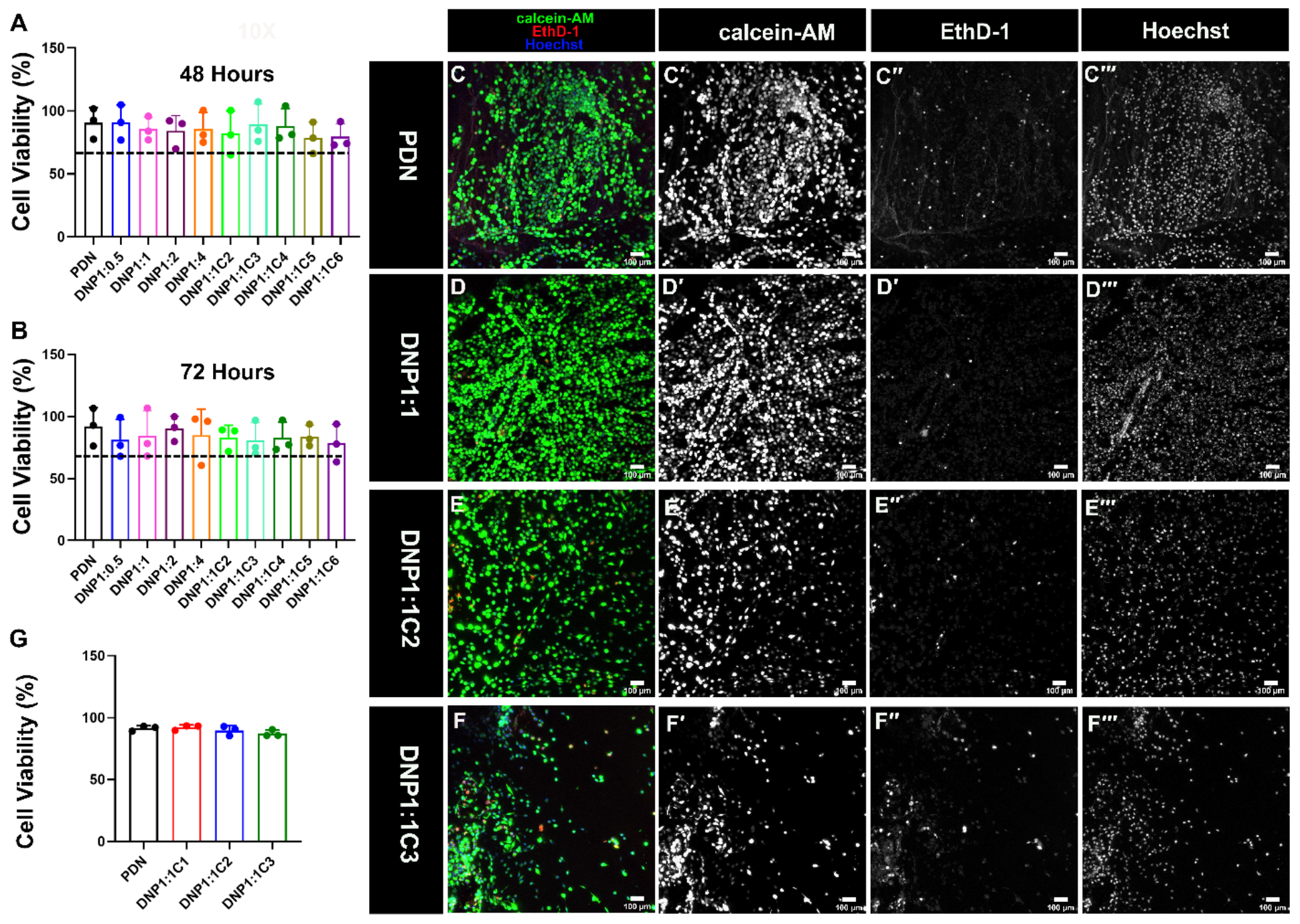
Biocompatability assay evaluating compatibility of biohybrid nerves with L929 (mouse fibroblast) cell lines. MTT assay reporting biocompatibility for all the samples at (A) 48 hours (B) 72 hours. The dotted line represent 70% cell viability, which is considered as a cutoff for the material to be considered biocompatible. (C,C′,C′′,C′′′- F,F′,F′′,F′′′) Immunofluorescent images of cells taken after live/dead assay, where calcein-AM (in green) represents live cells and ethidium homodimer (EthD-1) (in red) represents dead cells in contact with the (C,C′,C′′,C′′′) PDN (D,D′,D′′,D′′′) DNP1:1C1 (E,E′,E′′,E′′′) DNP1:1C2 and (F,F′,F′′,F′′′) DNP1:1C3. (G) quantification of Live/dead assay. The single filter images for respective dyes are shown for clarity. Scale bar = 100 μm.

Hemolysis testing was performed as it provides a simple and reliable method to evaluate the suitability of a material for use with blood, which is crucial for tissue engineering applications. The original nerve PDN and all the biohybrid nerves underwent hemolysis tests. All of the nerve samples showed minimal hemolysis, with a percentage below 10% **(Figure S4)**. According to ISO 10993-4 standards, a material that exhibits a hemolysis percentage below 10% is considered compatible with blood (hemocompatible), while a material with hemolysis below 5% is considered highly hemocompatible[56, 57]. The biohybrid nerve (DNP1:1C2) of interest that showed excellent material properties had a hemolysis percentage of 4.79%, which falls within the highly hemocompatible range.

In summary, a comprehensive evaluation of the cytocompatibility and hemocompatibility of the engineered biohybrid nerves indicates that DNP1:1C2 exhibits a high level of compatibility with both cells (cytocompatibility) and blood (hemocompatibility). As a result, it shows great potential as a promising candidate for further investigation into its capability to promote the ingrowth of motor axons.

### 2.5. Immunohistochemistry to evaluate motor axons in-growths

The goal of engineering biohybrid nerves is to create a medium that supports the growth of motor axons, specifically to reconstruct axonal tracts in the future. To evaluate the potential of a biohybrid nerve, DNP1:1C2, in promoting such motor axon ingrowth, we generated hSCS using HuES3 Hb9:GFP reporter cell lines **(Figure 5A)** and co-cultured them with the biohybrid nerve DNP1:1C2 for 14 days. Both the biohybrid nerves and hSCS were co-cultured in a custom-made 3D PDMS mold to ensure close proximity throughout the experiment **(Figure 5B)**. This setup prevented any disruption or dislodgment of the motor axon projections from the hSCS into the nerve samples, avoiding any gaps or displacement. For comparison, we also co-cultured the untreated nerve (PDN) with hSCS using a similar 3D PDMS mold design **(Figure 5C,D)**.

**Figure 5.**
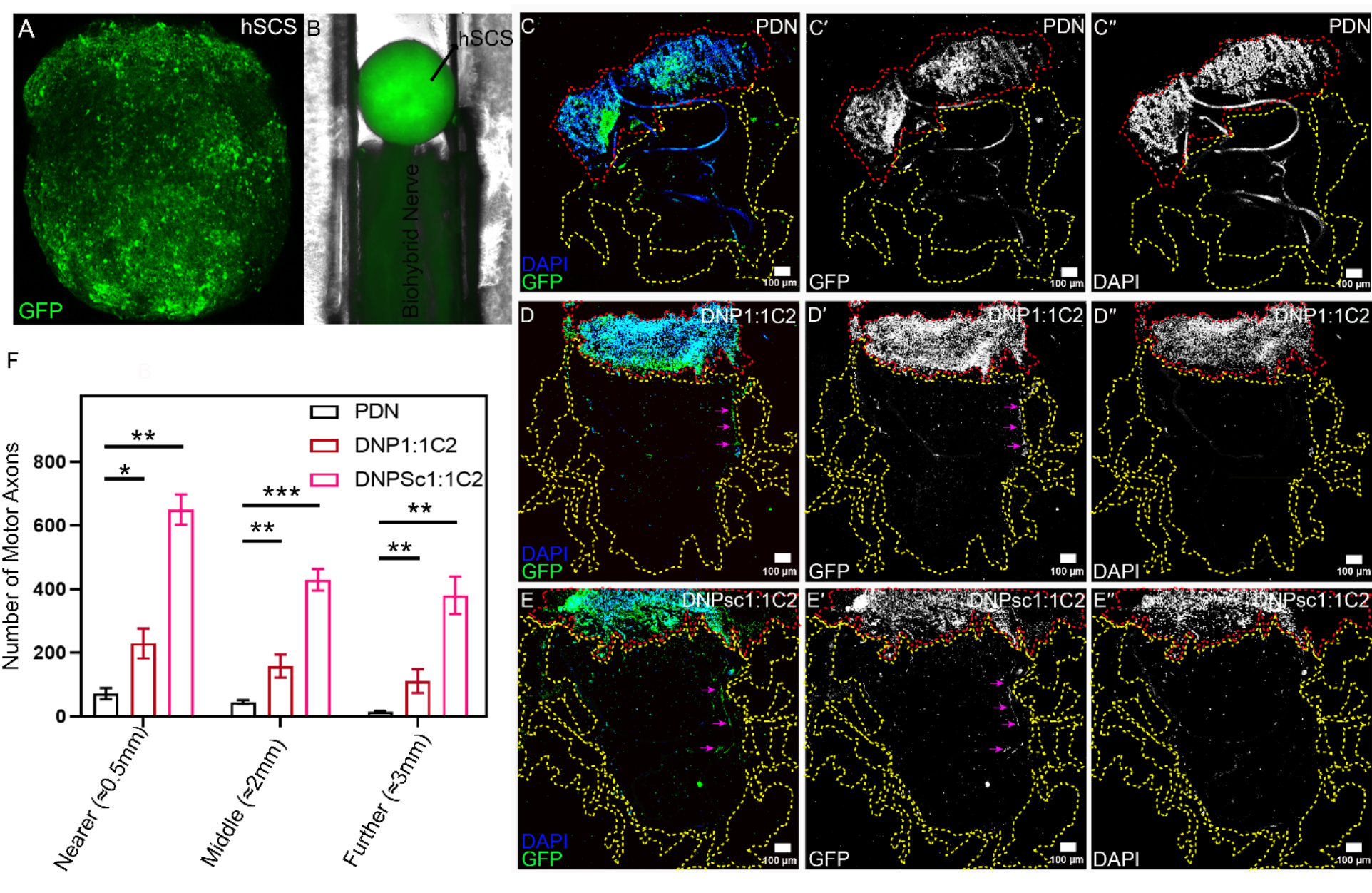
Motor axon in-growth projections get enhanced by biohybrid nerves. (A) GFP labelled hSCS (HUES 3 Hb9: GFP) (B) Experimental setup showing hSCS in contact with the biohybrid nerve placed together into the 3D printed PDMS mold. hSCS were cultured with PDN and Biohybrid nerves (DNP1:1C2 and DNPsc1:1C2) for 14 days followed by sectioning and (C,C′,C′′-E,E′,E′′) staining to image ingrowth of GFP labeled motor axons into the (C,C′,C′′) PDN and biohybrid nerve (D,D′,D′′) DNP1:1C2 and (E,E′,E′′) DNPsc1:1C2. The yellow dotted and red dotted line depicts boundary of nerve conduit and **hSCS** respectively. The single filter images for respective dyes are shown for more clarity. The GFP marker representing motor axons were calculated manually for each sample within the yellow dotted boundary. (F) The data clearly depicts that motor axons ingrowth are promoted by the biohybrid nerve in all the sections (nearer, middle and further). The pink arrows in (D,D′, E E′) highlights that neuron exhibit a propensity to grow and interconnect, forming cohesive network. Please note (C,C′,C′′) nerves without PEDOT coating show autofluorescence at few spots which is visible in all the channels (GFP as well as DAPI). Quantification of motor axons is done for sections from three different experimental samples (n=3) represented as mean ± SD, *P < .05, **P < .01. Scale bar = 100μm.

After co-culture, the nerve samples and hSCS assemblies were transversely sectioned and stained with GFP booster and DAPI **(Figure 5C,C′,C′′-E,E′,E′′)**. The data revealed a 4-fold increase in motor axon projections into the biohybrid nerve DNP1:1C2 compared to the pristine nerve PDN **(Figure 5F)**. To further enhance motor axon ingrowth, we seeded the biohybrid nerve (DNP1:1C2) with Schwann cells (DNPsc1:1C2) and interestingly found a 11-fold increase. One can also note that motor neurons exhibit a propensity to grow and interconnect, forming a cohesive network in case of biohybrid nerve compared to PDN as depicted by pink arrows **(Figure 5D,D′-E,E′)**. Additionally, we quantified neurites at the middle and furthest ends (from the hSCS) of the biohybrid nerve transverse sections and found a significantly higher number of neurites for both biohybrid nerves (DNP1:1C2) and (DNPsc1:1C2) when compared to PDN **(Figure S5A, A′, A′′-F, F′, F′′)**. DNP1:1C2 showed a 3.5-fold increase at the middle section and a 7.5- fold increase for sections cut at the further end of the nerve conduit compared to PDN. Similarly, DNPsc1:1C2 showed a 9.5-fold and 26-fold increase for middle and further end sections, respectively **(Figure 5F)**.

Our data strongly and clearly suggest that we have developed a promising biohybrid nerve, DNP1:1C2, and DNPsc1:1C2, which facilitate spinal motor axon ingrowth across the nerve conduit and can serve as a unique method to provide a desired medium for axonal reconstruction.

## 3. Discussion

Our findings show that PEDOT has the ability to seamlessly and inconspicuously integrate within decellularized nerve tissue through *in situ* polymerization, thereby improving its bioelectronic cues, while maintaining biophysical and biochemical characteristics. Following the analysis of the material, we found that among the 9 tested biohybrid nerves, DNP1:1C2 exhibited excellent conductivity, low stiffness, preserved mesostructures, and high level of compatibility with cells and living tissue (blood). Importantly, it notably enhances motor axon projections in comparison to a decellularized autograft, PDN.

Recent advances in neural engineering technologies have led to the development of grafts aimed at reconstructing axonal tracts and repairing nerve gap injuries [9, 58–60]. Autografts or allografts are common routes to bridge the nerve gap, which otherwise cannot be repaired through conventional end-to-end surgeries [9, 61]. However, these approaches have certain limitations. Autografts may cause morbidity at the donor site, while allografts may face graft rejection due to a mismatch in the major histocompatibility complex (MHC) (alloimmunity) [9]. Alternatively, artificial neural grafts can be implanted within the nerve stump, where the damaged nerve’s distal and proximal ends are connected to such nerve guidance conduits to reconstruct axonal tracts. However, many of these conduits lack intrinsic electrical properties to stimulate the ingrowth of axons and migration of Schwann cells [62]. Ideally, the conduit should have minimal mechanical mismatch to prevent compression of regenerating axons and tissue ischemia. Its mesostructures should closely resemble those found in natural nerves, providing an optimal guidance channel for axons and Schwann cells to traverse. Most importantly, the conduit should be compatible with the host’s cells and blood, seamlessly integrating with the surrounding tissue and withstanding the physiological environment without triggering an immune response [63–67].

The biohybrid nerve conduit (DNP1:1C2) we have developed offers unique advantages in the context of reconstructing axonal tracts as it closely matches all the criteria designated for an ideal conduit. One of its key strengths is its electrical properties, as evidenced by its conductivity of 0.214 mS cm-1 **(Figure 2D)**. The conductivity of a nerve conduit directly influences cellular activities, such as axonal ingrowth [62, 68]. Previous studies have shown that as conductivity increases to 1.1 X 10^-3^ mS cm^-1^ and 7 X 10^-2^ mS cm^-1^, respectively, the cellular response of nerve conduits improves [69, 70]. However, some reports indicate that when conductivity is below 0.1 mS cm^-1^, the material fails to exhibit the desired biologically transformational effect [71].Our biohybrid nerve conduit’s conductivity surpasses what has been previously reported for nerve conduits and exceeds the threshold of 0.1 mS cm^-1^. This strongly suggests that the developed biohybrid nerve can provide significant electrical cues to initiate cellular activities, such as axonal ingrowth. Moreover, the improved conductivity of the biohybrid nerve, DNP1:1C2, is mainly attributed to the conductive polymer PEDOT and recent studies on electroactive tissues, including nerves, have demonstrated the success of variously doped PEDOT conductive materials in electrically stimulating cells to promote cellular adhesion [72], proliferation [73], migration [26], and regulate axonal outgrowths (neurite outgrowths)[74, 75]. Furthermore, it is important to note that introducing a conductive material into a decellularized nerve presents a tradeoff between electrical and mechanical properties [76]. However, in the case of our biohybrid nerve (DNP1:1C2), the stiffness did not undergo significant changes compared to the pristine autograft **(Figure 3B),** thus validating its suitability as a candidate for neuronal interface.. Additionally, our designed biohybrid nerve closely matches the internal architecture of an autograft, which can provide crucial scaffolding to support the migration of axons **(Figure 3)**. This eliminates the need to rely on the formation of a stable fibrin clot as a guiding structure to facilitate axonal migration, which is a limitation of many artificial nerve conduits [36, 77, 78].These artificial nerve conduits either fail to form a stable fibrin clot or do not closely resemble the nerve’s internal structure [36, 77, 78].The biohybrid nerve we designed also exhibits high compatibility with both cells and blood **(Figure 4 and Figure S4)**. The cytocompatibility is critical for success of any nerve conduit as cytotoxicity can induce cell death, inflammation, immunogenicity, cause graft versus host disease, degrade conduit thus impeding its performance to reconstruct axonal tracts [62, 79–81]. Similarly, hemocompatibility of a nerve conduit is important as nerve conduit can come in direct contact with the blood and should be hemocompatible to avoid triggering immune responses, blood clotting, hemolysis (rupturing of red blood cells), or other detrimental effects to the recipient [82, 83].

To validate the potential for axonal growth in the biohybrid nerves we developed a 3D model that represents human spinal motor neurons. In humans, spinal motor neurons are found in the ventral horn of the spinal cord and are distributed along the anterior-posterior axis across various spinal sections, including cervical, thoracic, lumbar, and sacral regions [84, 85].The hSCS developed in this study can simulate the conditions in which spinal motor axons extend into the nerve scaffold, resembling the in vivo environment [86]. Additionally, this 3D model system offers advantages over animal models, as we can tailor the *in vivo* microenvironment as per the experimental purposes [87, 88]. In our study, we co-cultured biohybrid nerves with hSCS in a controlled environment within a custom-designed mold. The data clearly demonstrated that biohybrid nerves significantly enhance the growth of motor axons into the nerve scaffold compared to the pristine autografts **(Figure 5)**. This transportation of axons is likely initiated by PEDOT, which acts as an extracellular matrix filler, enhancing the biophysical and biochemical properties of a decellularized nerve. It is important to note that adding source of neurotropic factors like schwann cells [89, 90] further improved the results.

## 4. Conclusion

Our study has demonstrated the ability to create a biohybrid nerve by combining PEDOT with a decellularized nerve autograft, thereby harnessing the benefits of both components. The use of conductive polymer provides enhanced electrical properties, while the decellularized nerve graft maintains a structure closely resembling that of a natural nerve. Moreover, the process of decellularization reduces the likelihood of an immune reaction caused by a mismatch by MHC. The resulting biohybrid nerve is both compatible with cells and compatible with blood. Importantly, this hybrid characteristic significantly enhances motor axon ingrowth compared to the decellularized autograft alone, which ameliorates after seeding purified SCs. Looking ahead this technology has the potential to make a significant impact by enabling the reconstruction of axonal tracts in cases of severed nerve injuries, which is meaningful to the approximately 200,000 Americans who suffer from various types of peripheral nerve damage.

## 5. Materials and Methods

### 5.1. Harvesting and decellularizing Rat sciatic nerve

The sciatic nerves of male Sprague Dawley rats were obtained from fresh rat cadavers within 12 hours after death **(Figure 1A).** These were healthy rats without any known peripheral nerve damage. Rats were euthanized under carbon dioxide and aseptic techniques were used to isolate sciatic nerves which were then cleared of peripheral fat and connective tissue [91, 92] **(Figure 1B)**. Animal care and procedures were reviewed and approved by the Institutional Animal Care and Use Committee (IACUC) of Northwestern University, Protocol number: IS00014870. The decellularization process was carried out following the protocol previously described by Sondell et al [35].To initiate the decellularization process, the nerve samples were frozen at -20°C in sterile PBS (phosphate-buffered saline) and subsequently thawed. This freeze-thaw step was repeated three times to induce mechanical agitation, and each time the PBS was replaced with fresh PBS. Next, the nerves were treated with a series of detergents, including 3% Triton X-100 and 4% Sodium deoxycholate (SDC). These detergents were prepared in deionized (DI) water. The first step involved treating the nerve with Triton X-100 overnight, followed by three rinses with PBS. This treatment helped untangle and remove attached fat, further facilitating the cleaning process before the overnight treatment with SDC. The next day, the nerves were washed again with PBS three times to ensure removal of any debris or attached fat. The series of detergent treatments and PBS washes were repeated once more. Finally, the nerves were washed with DI water and stored in 10mM PBS (pH 7.2) at 4°C until further use **(Figure 1C)**.

### 5.2. Functionalizing nerves

The *in-situ* polymerization of EDOT (Sigma Aldrich #Cat: 483028-10G) was conducted using a method described previously [47, 93]. Initially, to remove any remaining detergent from the decellularized rat sciatic nerve constructs, they were subjected to sonication for 10 minutes in DI water with repeated rinses (three times), replacing the DI water each time. Next, the polymerization of EDOT took place on the rat sciatic nerve constructs using ferric chloride (FeCl3) (Thermo Scientific #Cat: 012357.09) oxidant in the presence of ethanol. The decellularized nerve (DN) constructs were conditioned at -20°C for 20 minutes in ethanol and EDOT, with a concentration that maintained the molar ratio of EDOT:FeCl3 at either 1M:0.5M (DNP1:0.5C1), 1M:1M (DNP1:1C1), 1M:2M (DNP1:2C1) or 1M:4M (DNP1:4C1). Subsequently, FeCl3 solution was added, thoroughly mixed, and kept at -20°C for 40 minutes to slow down the exotherm of polymerization process. The DN constructs were then left to air dry overnight in a fume hood, allowing the monomer to polymerize. The DN constructs with a molar ratio of EDOT: FeCl3 equal to 1M:1M were subjected to subsequent polymerizations 1 to 5 additional times providing 2 to 6 total polymerization cycles (DNP1:1C2, C3, C4, C5, C6). After each cyclic polymerization step, the DN constructs were sonicated for 10 minutes in deionized water, repeated three times, to remove any remaining Fe residue. Following the above method we generated 9 biohybrid nerve containing PEDOT:Cl as a conductive polymer. Throughout our manuscript we refer to it as PEDOT.

### 5.3. Electrochemical Impedance Spectroscopy

To assess the electrical properties and compare conductivity among pristine non-decellularized nerves (PnDN), pristine decellularized nerves (PDN), and biohybrid nerves, we performed Electrochemical Impedance Spectroscopy (EIS) using a PalmSens4 (version PSTrace 5.9.4515) potentiostat, as described in previous studies [94, 95]. The nerves used in the experiments were 3-5 mm in length with a 0.8-1 mm diameter. The nerves were first submerged in DI water overnight followed by 10 minutes sonication in DI water (repeated 5 times) to remove any residual ions from previously used electrolyte. To investigate the electric properties of the nerves, we employed a two-electrode system with a custom apparatus. The apparatus had gold coating on the edges of a glass substrate, with a 1 mm gap between two gold films. The design allowed the nerve to act as a bridge between the two gold electrodes, completing the electric circuit, and thus measuring changes in bulk resistance resulting from the decellularization process and various conditions in which decellularized nerves were modified with PEDOT. Furthermore, to enhance contact between the gold electrodes and nerve conduits, we applied a fixed weight affixed to the glass slide atop the nerve.

EIS measurements were performed from 0.1Hz to 10^6^ Hz, with a 10 mV RMS AC amplitude. After acquiring data we fitted data to the Kramers-Kronig (K-K) equation using AfterMath (1.6.10523) to validate if data has been acquired correctly. Any data that did not fit closely with the K-K (Chi^2^ value >0.05) was discarded. The data for which K-K fitting showed least error (Chi^2^ <0.05) was further fitted to an equivalent RC circuit model: R1(Q1||(R2Q2)) for PDN and PnDN (nerves that were not coated with PEDOT) and R1(Q1||R2)(Q2||R3) for biohybrid nerves. Here R1 represents uncompensated resistance, R2 and R3 denotes charge transfer resistance or bulk resistance (RCT), and Q1 and Q2 represents constant phase elements. For circuit fitting, we only considered frequencies between 1Hz and 10000 Hz. The values of R1, R2, R3, Q1 and Q2 were extracted based on the circuit fits. To calculate the conductivity of the nerves, we used the formula: S = T / (RCT × A), where S represents the conductivity, T and A are the thickness and area of the nerve, respectively, and RCT is the bulk resistance obtained after circuit fitting.

### 5.4. Mechanical Testing

Mechanical properties of nerve samples were analyzed via compression test using Piuma Nanoindenter (Optics11 life, Netherland) as described previously [96, 97]. Briefly, the lyophilized nerve samples were fixed to a petri dish with a double-sided Kapton tape and rehydrated with PBS overnight for equilibration. The ASTM F451-95 guidelines were followed to measure compressive mechanical strength of nerve samples using a probe that has glass spherical tip (9.5 µm in radius) mounted on a calibrated cantilever with stiffness of 0.039 N/m. A linear Hertzian contact model was used to calculate the effective Young’s modulus of the nerves [98].

### 5.5. Scanning Electron Microscopy (SEM) and Energy-dispersive X-ray Spectroscopy (EDS)

We utilized the Quanta 650F ESEM instrument to examine the morphology and microstructure of nerves under various conditions (PDN, DNP1:0.5C1, DNP1:1C1, DNP1:2C1, DNP1:4C1, DNP1:1C2, DNP1:1C3, DNP1:1C4, DNP1:1C5, DNP1:1C6).

Scanning Electron Microscopy (SEM) was employed to understand mesostructure, employing an accelerating voltage of 15 kV, spot size of 6 µm, and aperture of 3 µm. Before imaging, the nerve specimens (PDN) that lacked a conductive coating like PEDOT were coated with a 15 nm layer of Au/Pd using a Denton III Desk Sputter Coater. ImageJ software was used to measure the size of structural features from SEM micrographs. EDS mapping was conducted using the Quanta 650F ESEM instrument to visualize the distribution of Fe over the top surface of biohybrid nerves. The EDS maps of the specimens were analyzed using AZtec software.

### 5.6. X-ray Fluorescence

X-ray fluorescence spectroscopy was performed using Xenemetrix Ex-Calibur EX-2600 spectrometer equipped with an Rh X-ray tube and a silicon energy-dispersive detector. Each solid sample was placed in a plastic cup and powder was supported on a 6 μm Mylar film. Data collection was performed under vacuum and at room temperature, at 30 keV and 100 μA, and fluorescence spectra were collected for 60 s. No filter was used between the detector and sample and L-lines of Rh-radiation were visible in the collected spectra. Relative sulfur analysis was performed by integrating the fluorescence of the Kα1 transition for sulfur at ∼2.3 keV of each material. The residual Fe was quantified within each sample, using a calibration curve. Samples were compared under the same excitation settings and collected fluorescence was normalized by the mass of each sample.

### 5.7. Fourier Transform InfraRed Spectroscopy (FTIR)

The nerve composite was characterized by infrared spectroscopy of the powder using FTIR spectroscopy, which was performed using a Thermo Scientific Nicolet IS10 spectrometer. Samples were lyophilized and homogenized into KBr pellets for analysis.

5.8. *In Vitro* Cell Cytotoxicity

The ISO 10993-5 protocol was used to conduct *in vitro* cytotoxicity tests. The tests involved evaluating the cytotoxic effects of the specimen on cells through two methods: a live-dead assay for direct contact with the specimen and an MTT assay using extracts from the specimen. Prior to the cytotoxicity experiments, the samples were washed with ethanol and phosphate-buffered saline (PBS) using agitation (thrice for 15 minutes each).

For the live-dead assay, the following procedure was followed [99–102]: L929 cells, derived from L-connective mouse tissue strain, were obtained from ATCC (Cat no. CCL- 1) and seeded onto nerve samples using a dynamic bioreactor, as previously described [103]. Briefly, the nerve samples were combined with L929 cells in a 1.5 mL Eppendorf tube (3 nerve samples per tube) containing 1 mL of Dulbecco’s modified Eagle’s medium (DMEM)-high glucose media (Corning; Cat no. 10-013-CV) supplemented with 10% fetal bovine serum (FBS) and 1% antibiotic-antimycotic. The tubes were secured on a bioreactor rotator system with a rotation axis of 30° and a fixed speed of 18 rpm. The rotating device was placed in an incubator at 37 °C and 5% CO2 for 12 hours. After incubation, the seeded nerves were transferred to a culture dish containing DMEM media, and the cell culture media were changed after 48 hours and a live-dead assay was performed. To perform the live-dead assay, a solution was prepared using 0.5 μL of calcein AM, 2.0 μL of ethidium homodimer (EthD-1), and 1.0 μL of Hoechst in 1 mL of media. Each nerve sample was stained with 400 μL of the live-dead solution, incubated for 30 minutes, and imaged after destaining. The experiment was repeated three times for each condition, and the cell viability percentage was calculated using Matlab R2021B, as previously described [22, 101].

For the MTT assay, the following procedure was followed [100, 104, 105]. The nerve samples were treated with DMEM media (0.2 g/mL) in a 0.5mL Eppendorf tube. A control group without nerve samples also had media added to the tube. The tubes were incubated at 37 °C with 5% CO2 in a humidified incubator for 24 hours, while being agitated on a shaker at 70 rpm/min to create extract media. Separately, a suspension of L929 cells was prepared at a concentration of 4 × 104 cells/mL, and 100 μL of the suspension was dispensed into a 96-well plate. The plate was incubated at 37 °C with 5% CO2 in a humidified incubator for 24 hours. After 24 hours, the culture media was removed from the wells and replaced with 50 μL of extract media from the nerve samples, topped up with 50 μL of fresh culture media. The control group received "extract media" from the tube without nerve samples. For the positive control, the media was replaced with 20% DMSO (prepared in culture media), which caused severe cell death. For the negative control, 100 μL of culture media was used without cells, and the blank contained 100 μL of DMSO without cells. After 48 and 72 hours, the extract media and culture media were replaced with fresh 100 μL of culture media in their respective wells to which 10 μL of 12 mM MTT was added. The plates were incubated in the dark environment for 4 h, followed by the addition of 100 μL of the SDS-HCL solution to each well and thorough mixing. Plates were then incubated at 37 °C for 8 h, mixed thoroughly, and the absorbance was measured at 570 nm using UV–vis spectroscopy. MTT assays were repeated in triplicate.

### 5.9. Hemocompatibility

12 mL of the blood samples were collected from three female rabbits and pooled together in anticoagulant vials. The protocol to perform the hemolysis assays was adapted from prior literature procedures [57, 106]. Briefly, nerve samples were added to 4 mL of 0.9% saline solution and equilibrated for 30 min at 37 °C. 200 μL of diluted blood (4 mL of blood diluted in 5 mL of 0.9% saline solution) was added to each tube. A negative control was prepared by adding 200 μL of diluted blood to 4 mL of 0.9% saline solution (0% hemolysis) and a positive control was prepared by adding 200 μL of diluted blood to 4 mL of DI water (100% hemolysis). Each sample was then incubated at 37 °C for 1 h, centrifuged at 1000 rpm for 5 min, and the absorbance of the supernatant was measured at 545 nm. All the hemolysis experiments were performed in triplicate.

### 5.10. Generation of Hb9:GFP expressing 3D human Spinal Cord Spheroids (hSCS)

HuES3 Hb9:GFP reporter cells [107] were transformed into hSCS following a previously described method [86]. To summarize, HuES3 cells were cultivated in 6-well plates until they reached 70% confluency. On day 0, the cells were detached using Ethylenediaminetetraacetic acid (EDTA) (Thermo Fisher, #Cat no:15575020) and then transferred to ultra-low adhesion 96-well plates at a density of 10,000 cells per well. The cells were cultured in a motor axon base medium (Table S1) supplemented with 3µM Chir-99021 (Tocris, #Cat no: 4423), 0.2µM LDN 193189 (DNSK International, #cat no: 1062368-24-4), and 40µM SB 431542 (DNSK International, #cat no: 301836-41-9). On day 2, the cells were nourished with the base medium containing 3µM Chir-99021, 0.2µM LDN-193189, 40µM SB-431542, 100nM all-trans retinoic acid (RA), and 500nM smoothened agonist (SAG). hSCS were subsequently fed on days 4 and 7 with the base medium supplemented with RA and SAG. On day 9, the base medium was supplemented with RA, SAG, and DAPT. Finally, on day 11, the base medium was supplemented with RA, SAG, DAPT, Brain derived neurotropic factor (BDNF), and glial cell line derived neurotropic factor (GDNF). The maturation process occurred between days 14 and 35, during which the cultures were fed with hSCS maturation medium (Table S1) three times per week.

### 5.11. Seeding decellularized nerve with Schwann cells

A total of three biohybrid nerves (DNP1:1C2) were cultured with clinical-grade human SCs (sNF96.2) using a dynamic bioreactor, as previously described [100]. Biohybrid nerves were combined with SCs in a 15 mL tube containing 5 mL of cell culture medium and 1 million cells. The tubes were securely placed on a bioreactor rotator system with a rotation axis of 30° at a fixed speed. The rotating device was positioned in an incubator for 24 hours at 37°C and 5% CO2. Following incubation, the seeded biohybrid nerves were co-cultured with hSCS.

### 5.12. Co-culture of 3D human Spinal Cord Spheroids (hSCS) with biohybrid nerves

The experiment involved using 3D printed PDMS molds with a slot measuring 1.5mm wide, 2mm deep, and 5mm long. The slot was used to place biohybrid nerves (DNP1:1C2 and DNPsc1:1C2) in close proximity with hSCS. The PDMS molds were sterilized through autoclaving, while the biohybrid nerves underwent sterilization with 70% ethanol washes (15 minutes, repeated five times) before the experiment began. The biohybrid nerves were initially placed in the PDMS molds using sterile forceps. Subsequently, the PDMS molds containing the biohybrid nerves were transferred to 12-well cell culture plates. Mature hSCS were positioned in close contact with one end of the nerves using wide bore micropipette tips with a volume of 200 µL. The nerves and hSCS were then cultured together for a duration of 14 days. During this period, the hSCS medium (Table 1) was replaced every other day until the collection and fixation for immunohistochemistry.

### 5.13. Immunohistochemical analysis of Motor axon ingrowths

Nerve conduits along with hSCS were fixed with 4% paraformaldehyde and immersed stepwise in up to 30% sucrose/DPBS solution until they remained submerged. Following fixation and cryopreservation, samples were embedded in Optimal cutting temperature compound (OCT) and cryo-sectioned (5 μm thickness) for immunohistochemical analysis as described previously [108, 109]. Sections were blocked with 10% donkey serum in 0.03% Triton X-100/PBS and incubated with FITC Anti-GFP antibody (Abcam, Cat #ab6662). Nuclei were counterstained with DAPI (Fisher (Invitrogen), Cat #D1306). VECTASHIELD® Antifade mounting Medium (Vector Laboratories, Cat # H-1000) was used to mount coverslips, and slides were imaged with a Nikon AXR Confocal Laser Scanning Microscope as described previously [110, 111].

## Data Availability

The authors declare that the data supporting the findings of this study are available within the paper and its Supporting Information files.

## Acknowledgements

A.S.M, C.K.F. and J.R. acknowledge funding from NSF ASCENT (NSF ECCS 2023849). J.T., R.D., and J.R. acknowledge support from the Office of Naval Research (ONR) Young Investigator Program (YIP) Award No. N00014-20-1-2777. C.K.F. acknowledges funding from the Belle Carnell Regenerative Neurorehabilitation. This work utilized the Analytical bioNanoTechnology Equipment Core (ANTEC) and the Center for Advanced Microscopy/Nikon Imaging Center (CAM). Imaging work was also performed at the Northwestern University Center for Advanced Microscopy generously supported by CCSG P30 CA060553 awarded to the Robert H. Lurie Comprehensive Cancer Center. Our work also utilized the Keck-II facility of Northwestern University’s NUANCE Center and the Northwestern University Micro/Nano Fabrication Facility (NUFAB), both of which receive partial support from the Soft and Hybrid Nanotechnology Experimental (SHyNE) Resource (NSF ECCS-2025633), the Materials Research Science and Engineering Center (NSF DMR-2308691), the State of Illinois, and Northwestern University. Additionally, the Keck-II facility is partially supported by the International Institute for Nanotechnology (IIN), the Keck Foundation, and the State of Illinois, through the IIN.

