## Supplementary Information for "Decellularized biohybrid nerve promotes motor axon projections"

[a] Department of Biomedical Engineering, Northwestern University, Evanston, Illinois 60208, USA.
[b] Biologics Laboratory, Shirley Ryan Ability Lab, Chicago, IL, 60611 USA
[c] Division of Plastic Surgery, Feinberg School of Medicine, Northwestern University, 420 E Superior St., Chicago, IL, 60611 USA
[d] Section for Injury Repair and Regeneration Research, Stanley Manne Children's Research Institute, Ann & Robert H. Lurie Children's Hospital of Chicago, Chicago, IL 60611, USA
[e] Department of Pediatrics, Division of Critical Care, Northwestern University Feinberg School of Medicine, Chicago, IL 60611, USA
[f] Physical Medicine and Rehabilitation, Northwestern University Feinberg School of Medicine, Chicago, IL, 60611 USA
[g] Ken & Ruth Davee Department of Neurology, Northwestern University Feinberg School of Medicine, Chicago, IL, 60611 USA
[h] Department of Materials Science & Engineering, Northwestern University, Evanston IL. 60208, USA.
*

**Supplementary Table**

**Table S1: Ingredients of culture media needed to develop Hb9:GFP expressing 3D ventral Spinal Spheroids (vSO)**

| **ventral Spinal Spheroids (vSO) base media** | | |
|---|---|---|
| **Reagent** | **Vendor** | **Catalog number** |
| DMEM/F12 | Thermo Fisher | 11320-082 |
| Neurobasal | Thermo Fisher | 21103-049 |
| B27 Supplement | Thermo Fisher | 17504001 |
| N2 Supplement | Thermo Fisher | 17502001 |
| Antibiotic/Antimycotic<br>β-mercaptoethanol | | |
| Laminin | Thermo Fisher | 23017-015 |
| Ascorbic Acid | Sigma-Aldrich | A4403-100MG |
| Y-27632 Rock Inhibitor | DNSK International | 1062368-24-4 |
| **ventral Spinal Spheroids (vSO) maturation medium** | | |
| DMEM/F12 | Thermo Fisher | 11320-082 |
| Neurobasal | Thermo Fisher | 21103-049 |
| B27 Supplement | Thermo Fisher | 17504001 |
| N2 Supplement | Thermo Fisher | 17502001 |
| Antibiotic/Antimycotic | | |
| β-mercaptoethanol | | |
| Laminin | Thermo Fisher | 23017-015 |
| Ascorbic Acid | Sigma-Aldrich | A4403-100MG |
| Y-27632 Rock Inhibitor | DNSK International | 1062368-24-4 |
| Retinoic Acid (RA) | Sigma-Aldrich | Sigma Aldrich |
| GDNF | R&D Systems | 212-GD |

| | | |
|---|---|---|
| BDNF | R&D Systems | 248-BD |
| Smoothened Agonist (SAG) | DNSK International | 364590-63-6 |
| DAPT | Tocris | 2634/10 |

**Table S2: Circuit fit values for nerve samples.**

| Sample | Circuit Fit | R1 | R2 | R3 | $R_{CT}$ | $Chi^2$Value |
|---|---|---|---|---|---|---|
| PDN | R1(Q1p(R2sQ2)) | 34296.6898 | 124729.5986 | n/a | 124729.5986 | |
| PnDN | R1(Q1p(R2sQ2)) | 1.439059 | 26462.85253 | n/a | 26462.85253 | 0.05 |
| DNP1:1 C1 | R1s(Q1pR2)s(Q2pR3) | 835.5366733 | 13052.32044 | 585.2200553 | 13637.5405 | 0.005 |
| DNP1:1 C2 | R1s(Q1pR2)s(Q2pR3) | 369.4607427 | 3576.152727 | 718.486732 | 4294.639459 | 0.005 |
| DNP1:1 C3 | R1s(Q1pR2)s(Q2pR3) | 295.6075457 | 3472.935214 | 568.7712457 | 4041.70646 | 0.008 |
| DNP1:1 C4 | R1s(Q1pR2)s(Q2pR3) | 126.1101397 | 1226.394648 | 220.6094327 | 1447.004081 | 0.0002 |
| DNP1:1 C5 | R1s(Q1pR2)s(Q2pR3) | 198.7679517 | 1267.765134 | 339.9632347 | 1607.728369 | 0.0001 |
| DNP1:1 C6 | R1s(Q1pR2)s(Q2pR3) | 1004.928927 | 26560.01937 | 3539.800107 | 30099.81948 | 0.002 |

**Supplementary Figures**

**Figure S1**

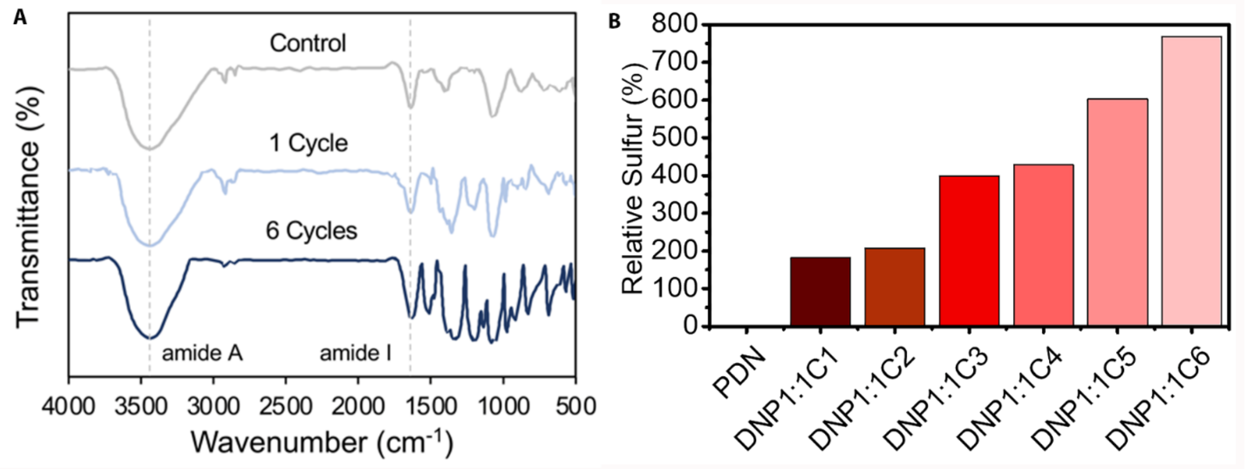


**Figure S1. The FTIR spectra displays characteristic bands for both collagen and PEDOT.** (A) The FTIR spectra of control nerve (PDN) (B) nerve sample representing minimum number (1 cycle) of cyclic polymerization with PEDOT (DNP1:1C1) and nerve sample representing maximum number (6 cycle) of cyclic polymerization with PEDOT (DNP1:1C6). The band at 3400 $cm^{-1}$ and 1625 $cm^{-1}$ are characteristic of collagen-based materials. Following the polymerization of EDOT, new bands emerge at around 1520 cm-1 and 1320 cm-1 corresponding to the stretching vibrations of C=C and C-C bonds in the thiophene ring and peaks at approximately 980 cm-1, 830 cm-1, and 680 cm-1 correspond to the stretching vibrations of C-S-C bonds. The XRF reports the (B) relative sulfur after polymerization of nerve samples with PEDOT to PDN.

**Figure S2**

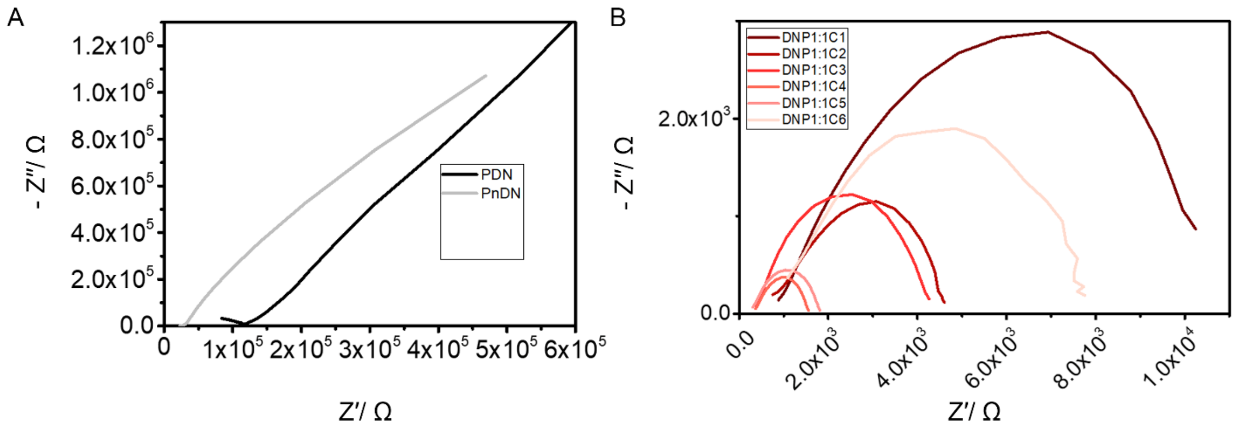


**Figure S2. Evaluating electrical properties of Biohybrid nerves.** The nyquist plot obtained from EIS characterization illustrators the relation between real impedance (Z′) and – imaginary impedance (Z″) for (A) pristine decellularized nerve (PDN) and pristine non-decellularized nerve (PnDN); and (B) biohybrid nerves with different cycles of polymerization (DNP1:1C1; DNP1:1C2; DNP1:1C3; DNP1:1C4; DNP1:1C5; DNP1:1C6). The shape of the Nyquist plot suggests both resistance and constant phase element.

**Figure S3**

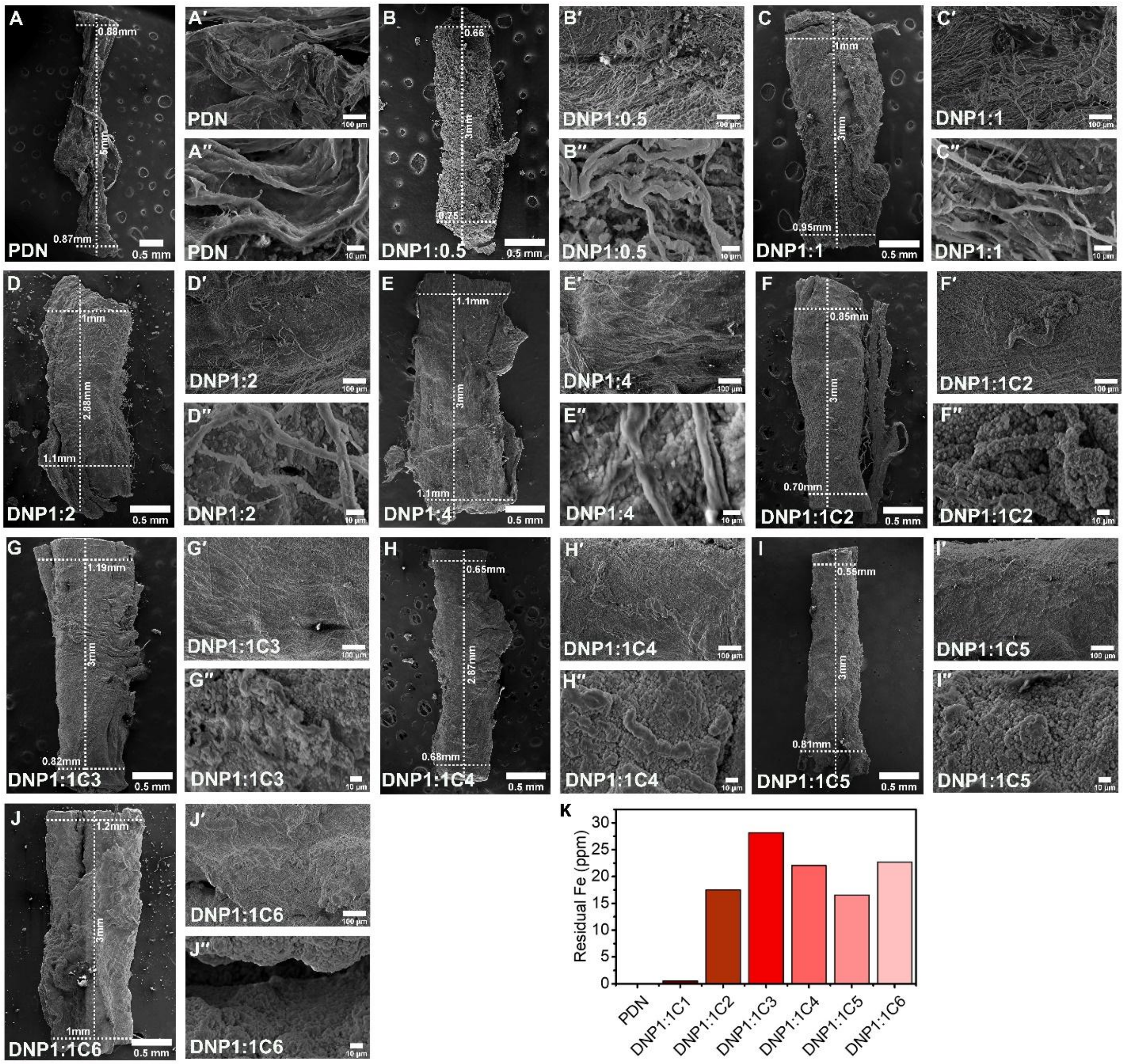


**Figure S3: Evaluation of Nerve Topology after *in-situ* polymerization with PEDOT.** Scanning electron microscopy images of biohybrid nerves (A,A',A'') PDN (B,B',B'') DNP1:0.5C1 (C,C',C'') DNP1:1C1 (D,D',D'') DNP1:2C1 (E,E',E'') DNP1:4C1 (F,F',F'') DNP1:1C2 (G,G',G'') DNP1:1C3 (H,H',H'') DNP1:1C4 (I,I',I'') DNP1:1C5 (J,J',J'') DNP1:1C6. Scale bar = 0.5 mm, 100 μm and 10 μm. The XRF reports the (K) residual Fe content.

**Figure S4**

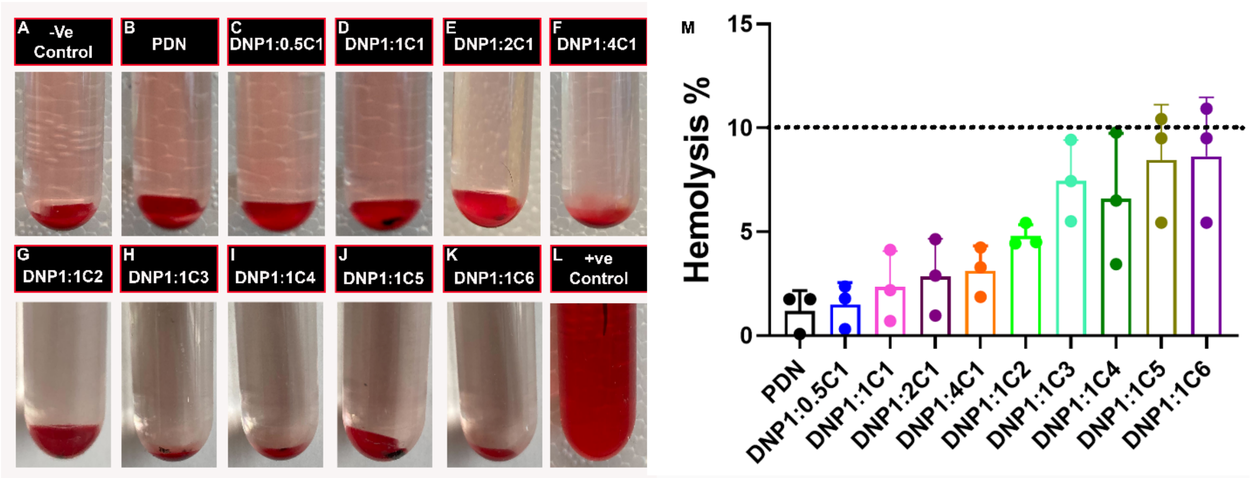


**Figure S4. Hemocompatability assay evaluating compatibility of biohybrid nerves with mammalian blood**. (A-L) Images of representative sample after incubation demonstrating a (A-K) lack of hemolytic reaction and hemolysis for (L) positive control. (M) Percentage hemolysis of pristine nerve (PDN) and biohybrid nerves. Each graph represents the mean ± SD ($n$ = 3). All the tested nerve samples show hemolysis less than 10 percent represented by the dotted line.

**Figure S5**

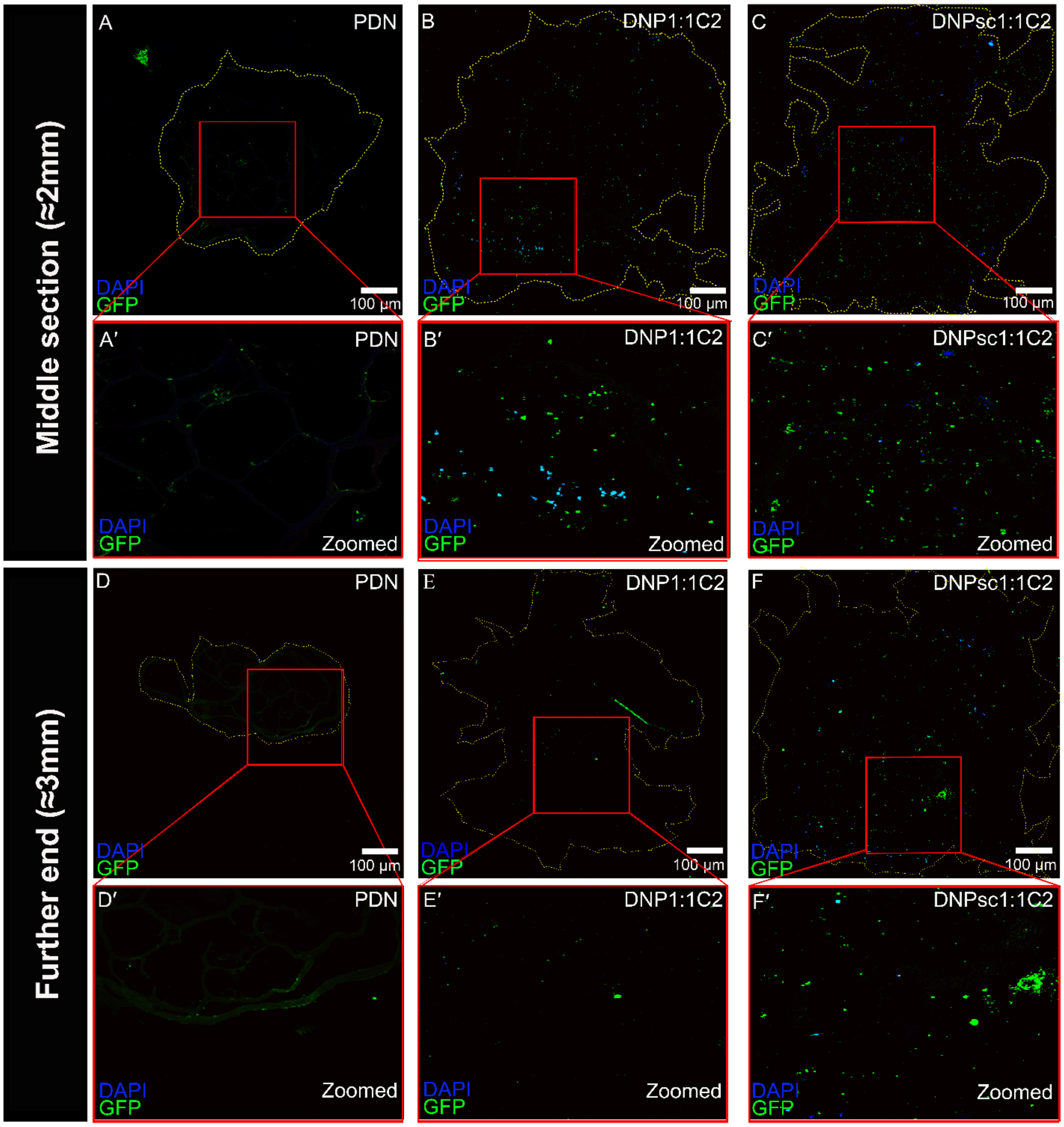


**Figure S5. Motor axon in-growth projects across the biohybrid nerves**. hSCS were cultured with PDN and Biohybrid nerves (DNP1:1C2 and DNPsc1:1C2) for 14 days followed by sectioning and (A,A′-F,F′) staining to image ingrowth of GFP labeled motor axons upto the middle section of (A,A′,) PDN (B,B′) DNP1:1C2 and (C,C′,) DNPsc1:1C2. Further end of (D,D′,) PDN (E,E′) DNP1:1C2 and (F,F′) DNPsc1:1C2. The yellow dotted line depicts boundary of nerve conduit. The zoomed images (A′-F′) for respective samples are shown for more clarity. The GFP marker representing motor axons were calculated manually for each sample within the yellow dotted boundary. The data clearly depicts that motor axons ingrowth are promoted across the biohybrid nerve. Quantification of motor axons is done for sections from three different experimental samples (n=3) represented as mean ± SD, in Figure 5 F.*P < .05, **P < .01, ***P < .001. Scale bar = 100 μm.
